# DataPackageR: Reproducible data preprocessing, standardization and sharing using R/Bioconductor for collaborative data analysis: A Practical Approach to Wrangling Multi-Assay Data Sets for Standardized and Reproducible Data Analysis

**DOI:** 10.1101/342907

**Authors:** Greg Finak, Bryan T. Mayer, William Fulp, Paul Obrecht, Alicia Sato, Eva Chung, Drienna Holman, Raphael Gottardo

## Abstract

A central tenet of reproducible research is that scientific results are published along with the underlying data and software code necessary to reproduce and verify the findings. A host of tools and software have been released that facilitate such work-flows and scientific journals have increasingly demanded that code and primary data be made available with publications. There has been little practical advice on implementing reproducible research work-flows for large ‘omics’ or systems biology data sets used by teams of analysts working in collaboration. In such instances it is important to ensure all analysts use the same version of a data set for their analyses. Yet, instantiating relational databases and standard operating procedures can be unwieldy, with high “startup” costs and poor adherence to procedures when they deviate substantially from an analyst’s usual work-flow. Ideally a reproducible research work-flow should fit naturally into an individual’s existing work-flow, with minimal disruption. Here, we provide an overview of how we have leveraged popular open source tools, including Bioconductor, Rmarkdown, git version control, R, and specifically R’s package system combined with a new tool *DataPackageR,* to implement a lightweight reproducible research work-flow for preprocessing large data sets, suitable for sharing among small-to-medium sized teams of computational scientists. Our primary contribution is the *DataPackageR* tool, which decouples time-consuming data processing from data analysis while leaving a traceable record of how raw data is processed into analysis-ready data sets. The software ensures packaged data objects are properly documented and performs checksum verification of these along with basic package version management, and importantly, leaves a record of data processing code in the form of package vignettes. Our group has implemented this work-flow to manage, analyze and report on pre-clinical immunological trial data from multi-center, multi-assay studies for the past three years.

## Introduction

A central idea of reproducible research is that results are published along with underlying data and software code necessary to reproduce and verify the findings. Termed a *research compendium,* this idea has received significant attention in the literature [1–5].

Many software tools have since been developed to facilitate reproducible data analytics, and scientific journals have increasingly demanded that code and primary data be made publicly available with scientific publications[2, 6–28]. Tools like git and github, figshare, and rmarkdown are increasingly used by researchers to make code, figures, and data open, accessible and reproducible. Nonetheless, in the life sciences, practicing reproducible research with large data sets and complex processing pipelines continues to be challenging.

Data preprocessing, quality control (QC), data standardization, analysis, and reporting are tightly coupled in most discussions of reproducible research, and indeed, literate programming frameworks such as Sweave and Rmarkdown are designed around the idea that code, data, and research results are tightly integrated[2, 25]. Tools like Docker, a software container that virtualizes an operating system environment for distribution, have been used to ensure consistent versions of software and other dependencies are used for reproducible data analysis[27]. The use of R in combination with other publicly available tools has been proposed in the past to build reproducible research compendia [3, 29, 30]. Many existing tools already implement such ideas. The *workflowr* package (https://github.com/jdblischak/workflowr#quick-start) provides mechanisms to turn a data analysis project into a version-controlled, documented, website presenting the results. The *drake* package [31] is a general purpose work-flow manager that implements analytic “plans”, caching of intermediate data objects, and provides scalability, and provides tangible evidence of reproducibility by detecting when code, data and results are in sync.

However, tight coupling of preprocessing and analysis can be challenging for teams analyzing and integrating large volumes of diverse data, where different individuals in the team have different areas of expertise and may be responsible for processing different data sets from a larger study. These challenges are compounded when a processing pipeline is split across multiple teams. A primary problem in data science is the programmatic integration of software tools with dynamic data sources.

Here, we argue that data processing, QC, and analysis can be treated as modular components in a reproducible research pipeline. For some data types, it is already common practice to factor out the processing and QC from the data analysis. For RNA sequencing (RNASeq) data, for example, it is clearly impractical and time consuming to re-run monolithic code that performs alignment, QC, gene expression quantification, and analysis each time the downstream analysis is changed. Our goal is to ensure that downstream data analysis maintains dependencies on upstream raw data and processing but that the processed data can be efficiently distributed to users in an independent manner and updated when there are changes.

Here, we present how the the Vaccine Immunology Statistical Center (VISC) at the Fred Hutchinson Cancer Research Center has addressed this problem and implemented a reproducible research work-flow that scales to medium-sized collaborative teams by leveraging free and open source tools, including R, Bioconductor and git[22, 32].

## Methods

### Implementation

Our work-flow is built around the *DataPackageR* R package, which provides a framework for decoupling data preprocessing from data analysis, while maintaining traceability and data provenance[18].

*DataPackageR* builds upon the features already provided by the R package system. R packages provide a convenient mechanism for including documentation as part of the built-in help system, as well as long-form `vignettes`, as well as version information and distribution of the entire package. Importantly, R packages often include data stored as R objects, and some packages, particularly under BioConductor, are devoted solely to the distribution of data sets[22]. The accepted mechanism for such distribution is to store R objects as `rda` files in the `data` directory of the package source tree and to store the source code used to produce those data sets in `data-raw`. The *devtools* package provides some mechanisms to process the source code into stored data objects[33].

Data processing code provided by the user (in the form of Rmd files preferably, and R files optionally) is run and the results are automatically included as package vignettes, with output data sets (specified by the user) included as data objects in the package. Notably, this process, while apparently mirroring much of the existing R package build process, is disjointed from it, thereby allowing the decoupling of computationally long or expensive data processing from routine package building and installation. This allows *DataPackageR* to decouple data munging and tidying from data analysis while maintaining data provenance in the form of a vignette in the final package where the end-user can view and track how individual data sets have been processed. This is particularly useful for large or complex data that involve extensive preprocessing of primary or raw data (e.g. alignment of FASTQ files for RNASeq or gating of flow cytometry (FCM) data), and where computation may be prohibitively long or involve software dependencies not immediately available to the end-user.

*DataPackageR* implements these features on top of a variety of *tidyverse* tools including *devtools, roxygen2, rmarkdown, utils, yaml, purrr.* The complete list of package dependencies is in the package `DESCRIPTION` file.

### Package structure

To construct a data package using *DataPackageR,* the user invokes the `datapackage.skeleton()` API, which behaves like R’s *package.skeleton(),* creating the necessary directory structure with some modifications. A listing of the structure of *DataPackageR* skeleton package directory, with other associated files is shown in Listing 2. The `datapackage.skeletion` API takes several new arguments apart from the package *name.* First, *code_files* takes a vector of paths to *Rmd* or *R* scripts that perform the data processing. These are moved into the `data-raw` directory by *package.skeleton().* The argument *r_object_names* takes a vector of quoted R object names. These are objects that are to be stored in the final package and it is expected that they are created by the code in the *R* or *Rmd* files. These can be *tidy* data tables, or arbitrary R objects and data structures (e.g. *S4* objects) that will be consumed by the package end-user. Information about the processing scripts and data objects is stored in a configuration file named `datapackager.yml` in the package root directory and only used by the package build process. The scripts may read raw data from any location, but generally the package maintainer should place it in `inst/extdata` if file size is not prohibitive for distribution.

### The build_package API

Once code and data are in place, the `build_package()` API invokes the build process. This API is the only way to invoke the execution of code in `data-raw` to produce data sets stored in `data.` It is not invoked through R’s standard `R CMD build` or `R CMD INSTALL` APIs, thereby decoupling long and computationally intensive processing from the standard build process invoked by end-users. Upon invocation of `build_package()` the *R* and *Rmd* files specified in `datapackager.yml` will be compiled into *package vignettes* and moved into the `inst/doc` directory, data objects will be created and moved into `data`, data objects will be version tagged with their *checksum* and recorded in the *DATADIGEST* file in the package root, and a *roxygen* markup skeleton will be created for each data object in the package.

### YAML Configuration

The `datapackager. yml` configuration file in the package root controls the build process by specifying which `R` and `Rmd` files should be processed and which `named R objects` are expected to be included as data sets in the package. Listing 1 shows such a configuration file. The API for interacting with this file is outlined in Table 1.

**Table 1:** The API for interacting with a yaml config file used by DataPackageR allows the user to add and remove data objects and code files, toggle compilation of files, and read and write the configuration to the data package.

| API call | Property | return value |
| --- | --- | --- |
| yml_add_files | file | config object |
| yml_remove_files | file | config object |
| yml_add_objects | objects | config object |
| yml_remove_objects | objects | config object |
| yml_find | config file | config object |
| yml_write | config file | null |
| yml_enable_compile | enable | config object |
| yml_disable_compile | enable | config object |

#### Listing 1

The structure of the YAML configuration file used by DataPackageR to control compilation and inclusion of data objects in the package..

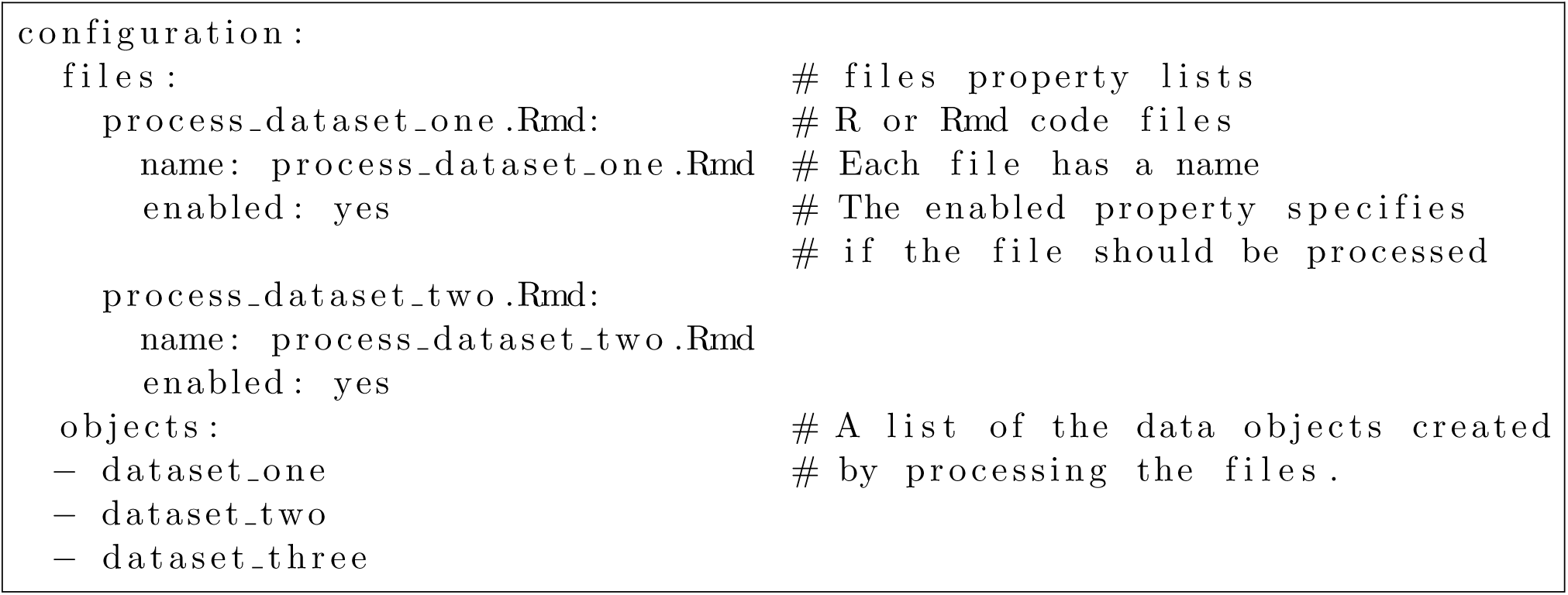

### Dataset Versioning

During the build, the `DATADIGEST` file is auto-generated. This file contains an `md5` hash of each data object stored in the package as well as an overall data set version string. These hashes are checked when the package is rebuilt; if they do not match, it indicates the format of the processed data has changed (either because the primary data has changed, or because the processing code has changed to update the data set). In these cases, the *DATADIGEST* for the changed object is updated and the minor version of the `DataVersion` string in the `DESCRIPTION` file is automatically incremented. The DataVersion for a package can be checked by the `dataVersion()` API, allowing end-users to produce reports based on the expected version of a data set (Figure 1).

**Figure 1:**
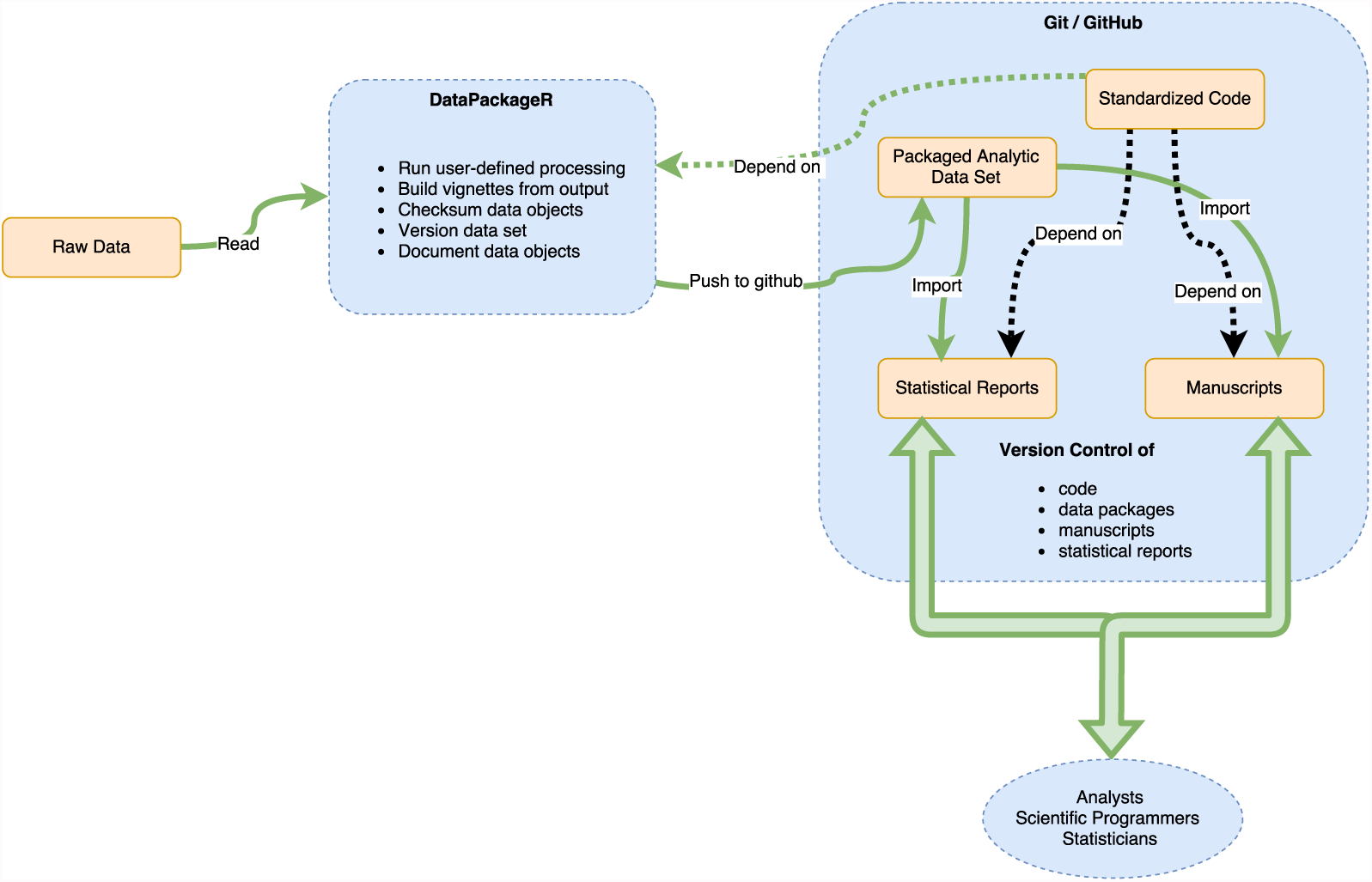
A schematic overview of the components in our reproducible data packaging work-flow that decouples data processing from data analysis.

#### Listing 2

The DataPackageR skeleton directory structure.

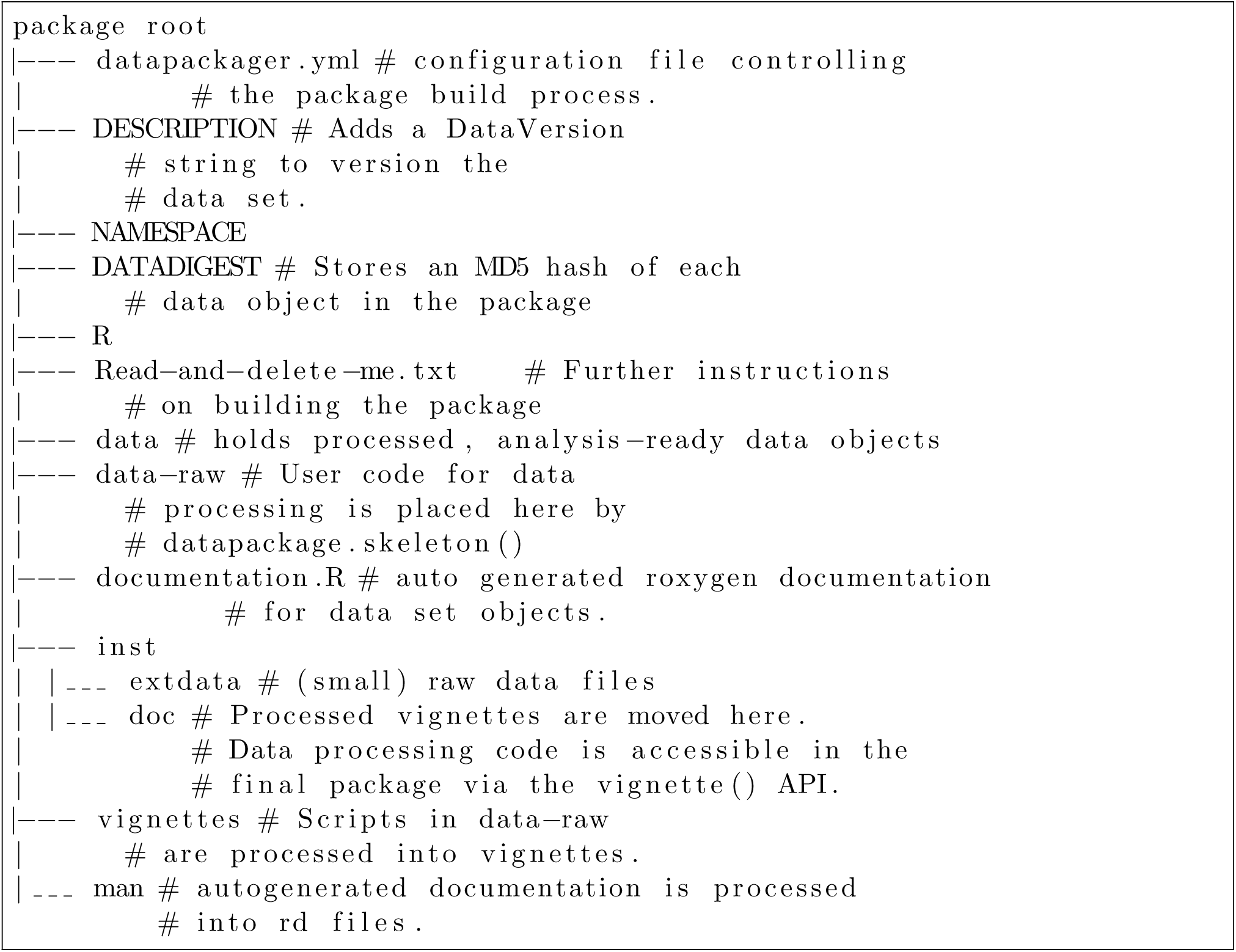

### Data Documentation

*DataPackageR* ensures that documentation is available for each data object included in a package by automatically creating an *roxygen* markup stub for each object that can then be filled in by the user. Undocumented objects are explicitly excluded from the final package.

Packages can be readily distributed in source or tarball form (together with the processed data sets under `/data/` and raw data sets under `/inst/extdata`). Within VISC we leverage *git* and *github* to provide version control of data package source code. By leveraging *DataPackageR,* the data processing is decoupled from the usual build process and does not need to be run by the end-user each time a package is downloaded and installed. Documentation in the form of *Rd* files, one for each data object in the package, as well as *html vignettes* describing the data processing, are included in the final package. These describe the data sets as well as how data was transformed, filtered, and otherwise processed from its raw state.

## Results

### Use Cases

*DataPackageR* was developed as a lightweight alternative to existing reproducible work-flow tools (e.g. *Galaxy[34]),* or to full fledged database solutions that are often beyond the scope of most short-term projects. *DataPackageR* plugs easily into any existing R-based data analysis work-flow, since existing data processing code needs only to be formatted into Rmarkdown (ideally). It is particularly suited for long-running or complex data processing tasks, or tasks that depend on large data sets that may not be available to the end user (e.g. FASTQ alignment or raw flow cytometry data processing). Such tasks do not fit well into the standard `R CMD build` paradigm, for example either as vignettes or *.R* files under `/data` since these would be invoked each time an end user builds a package from source. We desire, however, to maintain a link between the processed data sets and the processing code that generates them. We note that *DataPackageR* is distinct from other reproducible research frameworks such as *workflowr* or *drake* [31], in that it is designed to *reproducibly prepare data for analysis,* using an *existing code base,* with little additional effort. The product of *DataPackageR* is nothing more than an R package that can be used by anyone. The resulting data packages are meant to be shared, to serve as the basis for further analysis (Figure 1) and distributed as part of publications. These downstream analyses may leverage any of the existing work-flow management tools. Our goal is that data sets forming the basis of scientific findings can be confidently shared in their processed form which is often much smaller and easier to distribute.

Within the VISC, a team of analysts, statistical programmers, and data managers work collaboratively to analyze pre-clinical data arising from multiple trials. There are multiple assays per trial. The challenges associated with ensuring the entire team works from the same version of a frequently changing and dynamic data set, motivated the development of *DataPackageR.*

We demonstrate how *DataPackageR* is used to process and package multiple types of assay data from an animal trial of an experimental HIV vaccine.

### Data

We demonstrate the use of *DataPackageR* for processing data from a vaccine study, named *MX1,* designed to examine the antibody responses to heterologous N7 Env prime-boost immunization in macaques. The study had four treatment groups plus a control arm, with six animals per group. Samples were collected at three time points: t1: baseline, post-prime 2, post-boost 1, t2: post-boost 2, t3: post-boost 3. Six assays were run at each time point, using either serum samples or peripheral blood mononuclear cells (PBMCs). The assays were: 1) enzyme-linked immunosorbent assay (ELISA), an immunological assay that enables detection of antibodies, antigens, proteins and/or glycoproteins (serum); 2) a neutralizing antibody (Ab) assay (serum); 3) a binding antibody multiplex assay (BAMA) to assess antibody response breadth (serum); 4) a BAMA assay to permit epitope mapping (serum); 5) an antibody dependent cellular cytotoxicity (ADCC) assay (serum); and 6) an intracellular cytokine staining assay to assess cellular responses (PBMCs).

The raw data and environment to reproduce the processing *withDataPackageR* are distributed as a Docker image on `dockerhub.com` as `gfinak/DataPackageR:example`. We have restricted the number of FCS files distributed in the container to limit the size of the image and speed up processing of FCM data for demonstration purposes.

### Flow cytometry and other assay data

Flow cytometry (FCM) is a high content, high throughput assay for which VISC leverages specialized data processing and analytics tools. Raw FCS files and manual gate information in the form of FlowJo (FlowJo LLC, Ashland, OR) workspace files are uploaded directly to VISC by the labs. The raw data are processed with open source BioConductor software (*flow Workspace*) to import and reproduce the manual gating, extract cell subpopulation statistics, and access the single-cell event-level data required for downstream modeling of T-cell polyfunctionality and immunogenicity[35–37]. Tables of extracted cell populations, cell counts, proportions, and fluorescence intensities are included in study packages, together with an rmarkdown vignette describing the data processing. Due to the size of the raw FCS files, they are imported for processing from a location external to the data package source tree, so that the raw files are not part of the final package, but vignettes outlining the data processing are automatically included.

Remaining assay data are of reasonable size and are provided raw data in tabular (csv) form, imported into the package, processed and standardized from the `inst/extdata` package directory. Users can run and connect to an Rstudio instance in the container, where code and data to build the *MX1* data package reside.

## Summary

Reproducibility is increasingly emphasized for scientific publications. We describe a new utility R package, *DataPackageR* that serves to help automate and track the processing and standardization of diverse data sets into *analysis-ready data packages* that can be easily distributed for analysis and publication. *DataPackageR,* when paired with a version control system such as *git,* decouples data processing from data analysis while tracking changes to data sets, ensuring data objects are documented, and keeping a record of data processing pipelines as vignettes within the data package. The principle behind the tool is that it remains a lightweight and non-intrusive framework that easily plugs into most R-based data analytic work-flows. It places few restrictions on the user code therefore most existing scripts can be ported to use the package. The VISC has been using *DataPackageR* for a number of years to perform reproducible end-to-end analysis of animal trial data, and the package has been used to publicly share sets for a number of published manuscripts[36, 38, 39].

## Data availability

Partial data for the MX1 study to demonstrate processing using *DataPackageR* are available as a docker container from `gf inak/DataPackageR: latest`. The processed MX1 data are available on the CAVD DataSpace data sharing and discovery tool at http://dataspace.cavd.org under study identifier, CAVD 451.

### Software availability

This section will be generated by the Editorial Office before publication. Authors are asked to provide some initial information to assist the Editorial Office, as detailed below.

1. source code: (*http://github.com/RGLab/DataPackageR*)
2. reproducible examples: (*http://hub.docker.com/r/gfinak/datapackager/*)
3. License: MIT license

## Glossary

antibody: A protein produced by B cells that recognizes a specific antigen. Antibodies are released by B cells and bind to antigens that the body recognizes as foreign (non-self), such as bacterial and viral antigens. 7, 9
baseline: In vaccine trials this refers to a time point in the study before treatment is given, usually immediately before treatment. 7, 9
FASTQ: A standard, text-based file format for storing RNA and DNA sequence data along with quality information for individual nucleotide calls and some associated metadata. FASTQ files are the standard output of RNA sequencing experiments. 4, 9
FCM: Flow cytometry is a high content, high throughput assay enabling simultaneous multiparametric (i.e. cell-surface and intracellular protein abundance, or DNA content) measurement on suspended single-cells as they pass through the detection apparatus of a flow cytometer.. 4, 9
prime-boost: A vaccination strategy where one type of vaccine formulation is given to prime the immune system and another is given to boost the immune response. An example heterologous prime-boost modality is where a recombinant DNA formulation is given as the prime vaccine and a protein is given as the boost. 7, 9
QC: The process of checking data quality. 3, 9
RNASeq: Ribonucleic acid (RNA) sequencing technology used to measure gene expression in biological isolates from cells or tissues. 3, 4, 9
VISC: The Vaccine Immunology Statistical Center is a research team at the Fred Hutchinson Cancer Research Center working on analysis and integration of pre–clinical and clinical HIV vaccine trial data. 3, 7–10

## Acknowledgements

The authors wish to acknowledge the contributions of the members of the VISC and the CAVD Data Space (CDS) for contributions to testing and feedback on the software. We also acknowledge the Collaboration for AIDS Vaccine Discovery (CAVD), the Comprehensive Antibody Vaccine Immune Monitoring Consortium (CAVIMC) and the Comprehensive Cellular Vaccine Immune Monitoring Consortium (CCVIMC) as well as Dr. Shiu-Lok Hu.

